# A new universal pore measurement and clustering approach for surgical meshes

**DOI:** 10.1101/446450

**Authors:** Carolin Brune, Sebastian Vogt, Christian Peiper, Klaus Brinker, Jürgen Trzewik

## Abstract

The pore structure and pore size is a crucial characteristic of surgical meshes. A huge amount of different approaches for mesh classification based on pore size or materials are available. It is difficult to use these classification methods because of the large variety in knitting structures and pore shapes. No agreement on an established method to measure the pore size is available despite the fact that the surgical community agrees that pore size and geometry are crucial factors for the result of a hernia repair. In this publication a new approach to characterize meshes based on their pore shape and pore size is presented. The pore size is defined by the largest inscribed circle within a mesh pore and the smallest circumscribed circle outside the pore, in addition to hence calculated values. This allows a characterization with regards to tissue ingrowth and bridging behavior. The measurements are made using the scientific image processing software ImageJ and additional customized software plug-in. Since the program ImageJ is public domain and the plug-ins are available, the measurements can be reproduced. Furthermore, the presented pore descriptions can also be used for manual pore size verification.

## Introduction

The application of surgical meshes within the tension-free procedure is the method of choice for hernia repair [1]. A significant number of means of describing surgical meshes based on, for example, their physical or material properties already exist. However, there is no common definition of pore size and shape, hence no agreement on an established method to measure the pore size is available despite the fact that the surgical community agrees that pore size and geometry are crucial factors for the result of a hernia repair. In 1997, Amid classified four types of hernia meshes, where type I contains meshes with a pore size of more than 75μm. He mentioned that this is the minimum size to allow tissue ingrowth [2]. In 2007 Mühl et al. emphasized that it is also important to focus on the pore structures and pore sizes and less on the polymer type [3]. Mühl et al. introduced an image analysis system which measures the porosity of meshes and take into account contributions from pore structure and geometry [3]. Unfortunately, it is difficult to reproduce their results without access to the used software.

Another approach was described by Deeken et al. in 2011, who developed a pore size classification where pores were grouped in five classes, from microporous (<100μm) to very large (>2000μm) pores. However, the authors provide no detailed information about how these values were measured and do not focus on geometrical structures [4]. Since there is no uniformly valid definition for pore sizes it is difficult to compare results from different publications. The large amount of possibilities to describe pores and subsequent classifications lead to confusion and inconsistencies. There are controversial discussions on pore size based mesh classification methods and how to draw possible conclusions based on the classification results [5].

However, it is common agreement that the pore size is a crucial factor for the tissue ingrowth and mesh performance in situ and should therefore be taken in consideration for mesh characterization [6] [7] [8].

Prior to this approach our work group conducted a screening on pore size definitions within a group of surgeons (N=11) to figure out their view on pore geometrics and pore sizes. Based on the results of that research it can be shown that there is also no uniformity and general agreement among the surgeons on pore size definition, which reinforces the need for a consistent approach (see Figure 2).

Therefore, our work group came to the conclusion that it won’t be sufficient to characterize pores just on the largest distance between the fibers with regard to bridging behavior and tissue ingrowth. In addition, the pore shape should be evaluated to allow a characterization of pores in meshes using form factors. Furthermore, it has to be acknowledged that most meshes are designed with multiple pore shapes within one mesh. Those mesh constructions should be considered when describing the pore design of a mesh. One idea would be to split those pores in pore clusters which should be described separately. Therefore it is more promising to focus on pore characterization than on a classification of meshes.

In this publication we present a characterization method which defines the pore size in terms of the largest circle that fits inside a pore (largest distance to all fibers, considering the scar formation) and the smallest circle which fits outside the pore to describe the pore′s shape. Thus, different mesh and pore constructions can be taken into account and every mesh can be individually characterized. The ratio of the diameter of the largest inner circle and the diameter of the smallest outer circle can be used to distinguish between circular (ratio about 1) and narrow pores (ratio>1).

By means of PROLENE^®^ Soft Polypropylene Mesh (ETHICON), the use of this approach is shown.

This approach can be performed manually or by using a semi-automatic software algorithm, e.g. ImageJ plug-in.

## Materials and Methods

### Image capturing and pre-processing

For measuring the pore size of the textiles an image with an appropriate capturing device has to be taken. For this experimental evaluation a Keyence Microscope VR-3100 (Keyence Corporation, Osaka, Japan) is used. To ensure a correct scaling in a subsequent process step, the capturing device has to display a scale bar or a similar scale opportunity. A uniform background has to be chosen to reduce image noise. The magnification has to be sufficiently high to evaluate all fibres and the image field should be as big as possible to analyse a large number of pores to generate significant values. The program which is used for the image preparation as well as for the pore measurement itself is ImageJ. ImageJ is an image processing software, developed by the National Institute of Health in the United States of America, which is public domain and Java-based [9]. To extend the functions of ImageJ additional plugins can be programmed and added into the software program. Therefore, a customized ImageJ-plugin [10] with appropriate Image analysis algorithms [11] in has been developed by our group to characterize surgical mesh properties.

The image preparation takes place in four steps, including a scaling process, where pixels are transformed into a measureable unit, like mm. Afterwards the image has to be smoothed. Therefore, a median filter is used. The median filter considers all pixels which are within the chosen neighbourhood and replaces them by the median of all pixels. Subsequently, the colour threshold has to be adjusted. At this process step the uniform background and lighting is proved as an advantage. Because the plugin requires a binary image the image has to be transformed to a binary image.

### Pore measurements

This pore measurement is based on circles which fit exactly inside the pore (largest inscribed circle/ inner diameter = id) and circles which fits exactly outside the pore (smallest circumscribed circle /outer diameter = od). A pore specific form factor (ratio of id/od) is introduced to define the pore shape. A simple single-value diameter based pore size definition is not able to distinguish between a circular and a long narrow pore with the same diameter. Furthermore, the area based on the pixels inside the pore is calculated and the largest inner diameter and smallest outer diameter are displayed to characterize the pore.

### Algorithm and mathematical approach

#### Boundary of the pore

We used the particle analyser of ImageJ in order to define the inner outlined boundary, represented by black pixels which are besides at least one white pixel, as the boundary (orange line in figure in Table 1(a)).

**Table 1:**
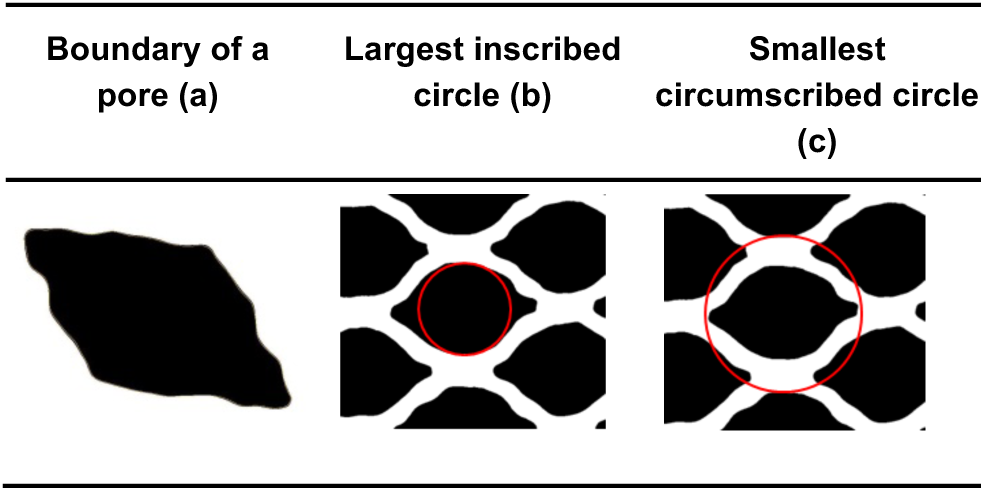
Visualization of boundary of a pore, largest inscribed circle and smallest circumscribed circle

#### Largest Inscribed Circle

The largest inscribed circle is the maximum circle which fits inside a pore. It is computed using the Euclidean Distance Map (EDM) provided by ImageJ. The algorithm is similar to the 8SSEDT of Leymarie and Levine [12].

The largest inscribed circle actually can be computed efficiently in ImageJ using the EDM by taking the pixel within a particle with the maximum value assigned. This pixel marks the centre and its value gives the radius (see figure in Table 1 (b)).

#### Smallest Circumscribing Circle

The smallest circumscribing circle is the minimal circle which fits around the pore. It is computed using only the convex hull of the outline of the pore [13] (see figure in Table 1 (c)).

#### Pore clustering

Because many surgical meshes contain a variety of different pores regarding pore shape and pore size, it is appropriate to cluster these pores in different types to avoid a distortion of the characterization values. This pore clustering process is a semi-automatic action. First, for each cluster a representative pore has to be determined. Afterwards, the plug-in calculates the matching pores and creates clusters (see Fig 1). Because of the high variety of pores and differences due to the knitting process, it is possible that some pores are classified wrongly. Hence the textile expert has the possibility to correct mistakes and reclassify pores.

**Fig. 1:**
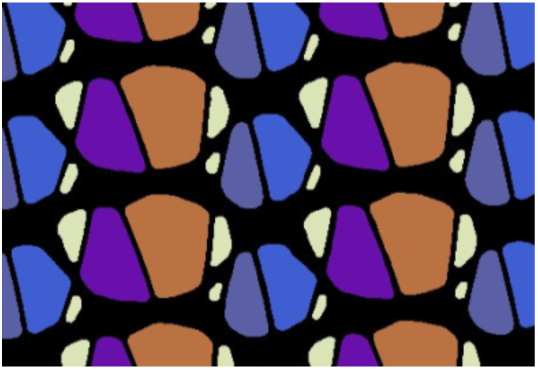
Pore clustering

#### Algorithm and mathematical approach

The pores can automatically be put in different categories, each category with similar pores. The categories are determined by the diameter of the largest inscribed circle, the diameter of the smallest circumscribing circle and the ratio of these, which is an indicator of the pore shape.

Before classifying each pore in a category, at least one pore has to be chosen manually to represent a category. Afterwards a nearest-neighbour algorithm is applied to find similar pores for each chosen category. To facilitate the data processing and comparison between pores, the categories are sorted on the basis of the diameter of the largest inscribed circle.

## Results

Obtained by a survey, the bar chart below (Figure 2) shows the variety of the surgeon’s ideas on the definition on a surgical mesh pore size. These widely spread answers depict the need of a uniform characterizing method for surgical meshes.

**Fig 2:**
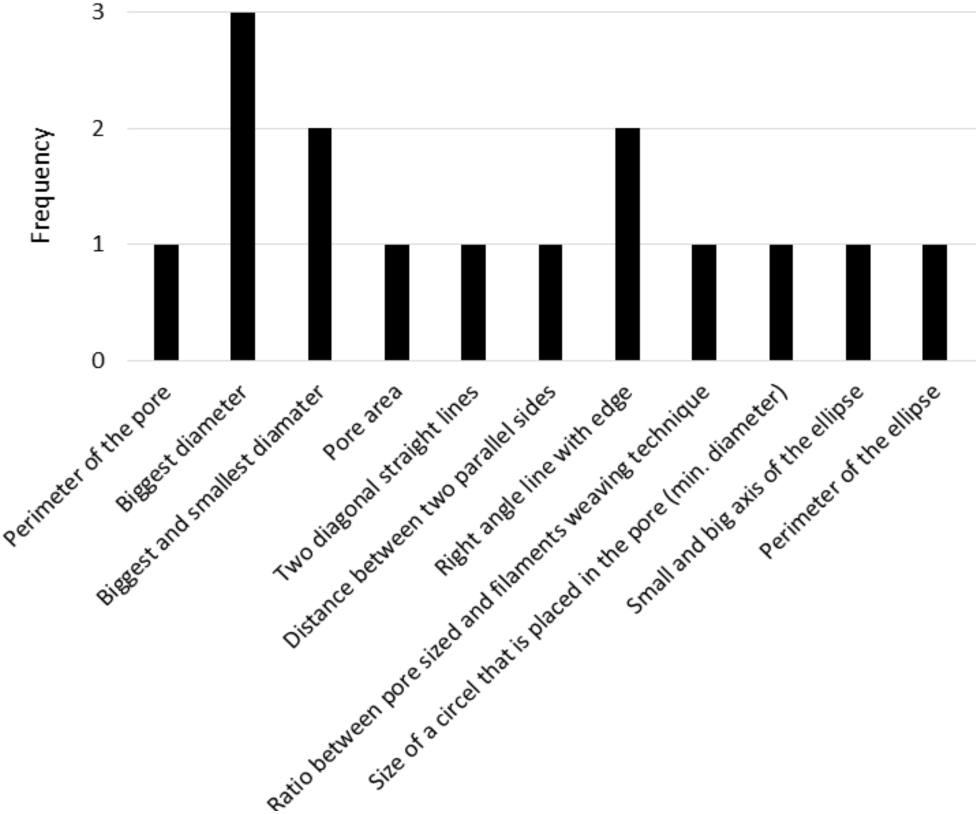
Survey based pore sizes definition by surgeons

The following data are calculated by the plug-in and displayed in a results table (see Table 2):

- Amount of pore types
- Ratio id/od
- Inner diameter [mm]
- Outer diameter [mm]
- Area [mm^2^]
- Percentage area [%]

**Table 2:** Results table

| Pore types | Mean Ratio id/od | Mean id [mm] | Mean od [mm] | Mean Area [mm<sup>2</sup>] | Percentage of total area [%] |
| --- | --- | --- | --- | --- | --- |
| 1 | 0,599 | 1,461 | 2,440 | 2,900 | 34,808 |
| 2 | 0,474 | 1,125 | 2,373 | 1,876 | 22,519 |
| 3 | 0,469 | 0,956 | 2,039 | 1,501 | 18,016 |
| 4 | 0,398 | 0,802 | 2,014 | 1,154 | 13,857 |
| 5 | 0,371 | 0,315 | 0,868 | 0,225 | 10,801 |

From this table, many conclusions regarding the mesh and pore structure can be made: the number of pore types provides information about the complexity of the mesh; the ratio id/od provides information about the shape of the pores (for the specific pore type); the inner diameter displays the size of the pore itself (with important information regarding tissue ingrowth and bridging behaviour). The outer diameter alone is less meaningful; however it is important to calculate the ratio. The area is a natural parameter in order to characterize the pore size; however it gives no further information about the pore shape. The percentage area is relevant to specify the pore type number in relation to all pores of the evaluated area. Some pores are defined as “remainder pores” (see white pores in Fig 1 and 3) these are pores which are too small to define them as significant pores. This has to be considered by the textile expert who conducts the measurement.

The following histogram shows the percentage of the single pore types within the complete pore area (Figure 3):

**Fig 3:**
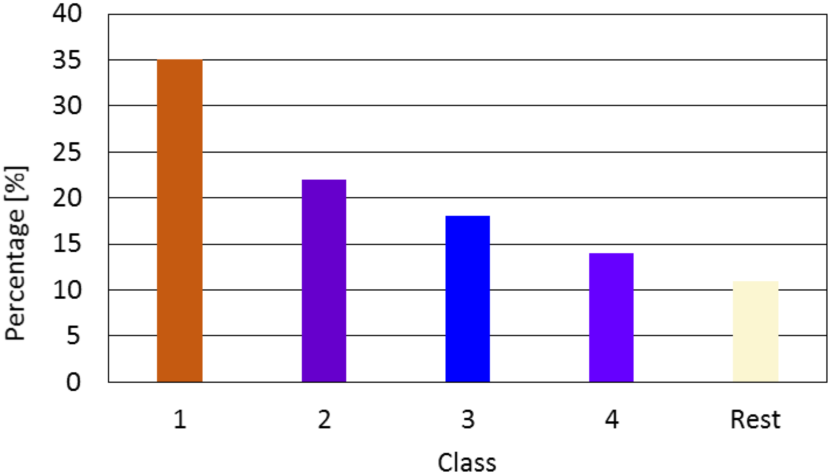
Histogram percentage of pore types

## Discussion

Because of the combination of an automatic and a semiautomatic approach, the program and the plug-in can apply to very complex and various different mesh types, all while following a prescribed, programmed approach. It is advised to characterize or even classify by not only one parameter, but by a manageable number of measured parameters.

Furthermore, this approach focuses on the geometric structure of the meshes and takes the important measured values for bridging and tissue ingrowth, which in particular is the inscribed circle or the smallest distance between fibres in all directions, in account.

This method provides a transparent and comprehensive clarification for surgeons and compares similar meshes without lumping all meshes together. Furthermore the presented pore descriptions can also be used for manual pore size verification.

Since the pore sizes vary under uniaxial tensile load in certain meshes [14], our method may help in objectivizing these differences. This may help to estimate or measure the porosity, if the mesh is implanted under mechanical strain, which occurs frequently in the TVT sling implant. This has a certain influence on tissue integration and foreign body response [15]. Meshes with adapted textile design promise stable porosity even under longitudinal or transverse strain [16], as it may occur in abdominal wall surgery or in hiatal reconstruction [17]. Our exact and reproducible possibility for measurement of these changes may help to answer these questions and to precisely define an effective porosity.

## Acknowledgment

We gratefully acknowledge the technical and textile support from Barbara Schuldt-Hempe and Boris Batke of Johnson & Johnson MEDICAL GmbH.

## Compliance with ethical standards

### Human and animal rights

This article does not contain any studies with human participants or animals performed by any of the authors.

### Informed consent

Informed consent was obtained from all individual participants included in the study.

## References

[1] J. Siewert, M. Rothmund and V. Schumpelick (2011) Gastroenterologische Chirurgie, 3. Ed., Springer Medizin Verlag, Heidelberg

[2] P. Amid (1997) Classification of biomaterials and their related complications in abdominal wall hernia surgery. Hernia, Vol. 1:pp. 15–21 doi: 10.1007/BF02426382

[3] T. Mühl, M. Binnenbösel, U. Klinge and T. Goedderz (2007) New objective measurement to characterize the porosity of textile impalants. Journal of Biomedical Materials Research Part B: Applied Biomaterials, Wiley InterScience, Vol. 84B: pp. 176-183, doi: 10.1002/jbm.b.30859

[4] C. R. Deeken, M. S. Abdo, M. M. Frisella and B. D. Matthews (2011) Physicomechanical Evaluation of Polypropylene, Polyester and Polytetrafluoroethylene Meshes for Inguinal Hernia Repair. Journal of the American College of Surgeons, Vol. 212: pp. 68–79, doi: 10.1016/j.jamcollsurg.2010.09.012

[5] A. C. d. Beaux (2012) Invited commentary on HERN-D-11-00301R3. Hernia, Vol. 16, no. 3: pp. S. 259–259, doi: 10.1007/s10029-012-0922-5

[6] A. M. Zihni, S. Ray, S. P. Lake, D. M. Thomson Jr., J. Gluckstein and C. R. Deeken (2014) Pore size and pore shape- but not mesh density-alter the mechanical strength of tissue ingrowth and host tissue response to synthetic mesh materials in a porcine model of ventral hernia repair. Journal of the mechanical behavior of biomedical materials, Vol. 42: pp. 186–197, doi: 10.1016/j.jmbbm.2014.11.011

[7] U. Klinge and B. Klosterhalfen (2012) Modified classification of surgical meshes for hernia repair based on the analyses of 1,000 explanted meshes. Hernia, Vol. 16: pp. 251–258, doi: 10.1007/s10029-012-0913-6.

[8] U. Klinge, J.-K. Park and B. Klosterhalfen (2013) The ideal mesh?. Pathobiology, Vol. 80: pp. 169–175, doi: 10.1159/000348446

[9] “ImageJ,” [Online]. Available: https://imagej.nih.gov/ij/. [Accessed 20 April 2016].

[10] “ImageJ plug-in” [Online]. Available: https://goo.gl/p0hWV0

[11] S. Vogt, C. Ritter, J. Trzewik, K. Brinker (2017) Measuring, clustering and classifying pores of surgical meshes with an ImageJ plug-in. 2nd YRA MedTech Symposium, Hochschule Ruhr West, June 8-9, Mühlheim a. d. Ruhr, Germany

[12] F. Leymarie and M. Levine (1992) A note on “Fast Raster Scan Distance Progatation on the Discrete Rectangular Lattice". CVGIP Image Understanding, pp. 84–94, doi: 10.1016/1049-9660(92)90008-Q

[13] J. Rokne (1992) An easy bounding circle. Graphic Gems II, pp. 14–16

[14] U. Klinge, J. Otto, T. Mühl, (2015) High structural stability of textile implants prevents pore collapse and preserves effective porosity at strain. BioMed Research International, Vol. 2015, pp. 1–7, doi: 10.1155/2015/953209

[15] W. R. Barone, P. A. Moalli, S. D. Abramowitch (2016) Textile properties of synthetic prolapse mesh in response to uniaxial loading. American Journal of Obstetrics and Gynecology, Vol. 215 (3), pp. 326–329, doi: 10.1016/j.ajog.2016.03.023

[16] A. Lambertz, R. R. Vogels, M. Binebösel, D.S. Schöb, K. Kossel, U. Klinge, U. P. Neumann, C. D. Klink (2015) Elastic mesh with themoplastic polyurethane filaments preserves effective porosity of textile implants. Journal of Biomedical Materials Research Part A, Vol. 103, pp. 2654–2660, doi: 10.1002/jbm.a.35411

[17] P. H. Alizai, S. Schmid, J. Otto, C. D. Klink, A. Roeth, J. Nolting, U. P. Neumann, U. Klinge, (2014) Biomechanical analysis of prosthetic mesh repair in a hiatal hernia model. Journal of Biomedical Materials Research Part B: Applied Biomaterials, Vol. 102: pp. 1485–1495: 10.1002/jbm.b.3312

